# Theory of Circadian Metabolism

**DOI:** 10.1101/321364

**Authors:** Michele Monti, David K. Lubesnky, Pieter Rein ten Wolde

## Abstract

Many organisms repartition their proteome in a circadian fashion in response to the daily nutrient changes in their environment. A striking example is provided by cyanobacteria, which perform photosynthesis during the day to fix carbon. These organisms not only face the challenge of rewiring their proteome every 12 hours, but also the necessity of storing the fixed carbon in the form of glycogen to fuel processes during the night. In this manuscript, we extend the framework developed by Hwa and coworkers (Scott *et al*., Science **330**, 1099 (2010)) for quantifying the relatinship between growth and proteome composition to circadian metabolism. We then apply this framework to investigate the circadian metabolism of the cyanobacterium *Cyanothece*, which not only fixes carbon during the day, but also nitrogen during the night, storing it in the polymer cyanophycin. Our analysis reveals that the need to store carbon and nitrogen tends to generate an extreme growth strategy, in which the cells predominantly grow during the day, as observed experimentally. This strategy maximizes the growth rate over 24 hours, and can be quantitatively understood by the bacterial growth laws. Our analysis also shows that the slow relaxation of the proteome, arising from the slow growth rate, puts a severe constraint on implementing this optimal strategy. Yet, the capacity to estimate the time of the day, enabled by the circadian clock, makes it possible to anticipate the daily changes in the environment and mount a response ahead of time. This significantly enhances the growth rate by counteracting the detrimental effects of the slow proteome relaxation.

## INTRODUCTION

Bacterial cells alter gene expression in response to nutrient changes in their environment [1–5]. In recent years, experiments have demonstrated that the relation between the proteome composition and the growth rate can be quantitatively described by growth laws, which are based on the idea that cells need to balance the supply of amino-acids via catabolic and anabolic reactions with the demand for amino-acids in the synthesis of proteins by ribosomes [1–5]. While in the original studies this relationship was tested for conditions that do not vary on the timescale of the cellular response [1–3], more recently it has been demonstrated that these growth laws can also describe the transient relaxation dynamics of the proteome in response to a nutrient shift [4, 5]. Here, we extend this framework to predict how bacterial cells repartition their proteome in response to periodic, circadian environmental changes.

Many organisms, ranging from cyanobacteria, to plants, insects, and mammals, possess a circadian clock, which means that they can anticipate daily changes in their environment, and adjust their proteome ahead of time. Moreover, many organisms face the challenge that they can fix carbon and/or nitrogen only during one part of the day, which means that they then need to store these resources to fuel processes the other part of the day. In this manuscript, we study by mathematical modeling the optimal strategy for allocating cellular resources that maximizes the growth rate of cyanobacterial cells living in a periodic environment. We show that storing carbon and nitrogen puts a fundamental constraint on the growth rate, and tends to generate extreme growth behavior, where cells predominantly grow in one part of the day. Moreover, we show that in cyanobacteria with cell-doubling times that are typically longer than 10h [6–8], the slow relaxation of the proteome severely limits the growth rate, but that anticipation makes it possible to alleviate the detrimental effects of the slow relaxation.

Cyanobacteria are among the most studied and best characterized organisms that exhibit circadian metabolism. Their metabolism is shaped by the constraint that not all the principal elements can be fixed during the day and the night. For cyanobacteria, the primary source of carbon is CO_2_, which they fix during the day via photosynthesis. Yet, cyanobacteria also need carbon during the night, not only for protein synthesis, but also for the generation of fuel molecules such as ATP, required for maintenance processes such as DNA repair. To this end, they use not all the fixed carbon to fuel growth during the day: they also store a fraction in the form of glycogen, which then becomes the principal source of carbon during the night.

Like all living cells, cyanobacteria not only need carbon, but also nitrogen. Some cyanobacteria, such as *Synechococcus* and *Synechocystis*, rely on nitrogen that has been fixed by other organisms in the form of, e.g., nitrate. Other cyanobacteria, such as *Cyanothece* [9] and *Anabaena* [10, 11], have, however, the ability to fix the nitrogen that is available in the form of the most abundant gas in the atmosphere, N_2_. Yet, this process requires an enzyme, nitrogenase, which cannot tolerate Ø2. Since Ø2 is produced during photosynthesis, cyanobacterial cells cannot simultaneously fix carbon and nitrogen during the day. *Anabaena* has solved this problem at the population level, where some cells fix carbon while others fix nitrogen [11]. *Cyanothece* has solved the problem at the single-cell level by temporally separating these processes [9]. In this manuscript, we will use *Cyanothece* as a model organism to study the design principles of circadian metabolism.

During the day *Cyanothece* stores carbon in the form of glycogen, while during the night it fixes nitrogen and stores it in the form of cyanophycin [12]. Like glycogen, cyanophycin is a large polymer that accumulates in the cytoplasm in the form of insoluble granules. The polymer is a large polypeptide and consists of two amino-acids: aspartic acid, which forms the back bone, and arginine, which constitutes the side group. Arginine is the amino acid with the largest number of nitrogen atoms in its side chain, namely 3; indeed, its side chain has the largest ratio of nitrogen (N) to carbon (C) atoms: 3:4. Cyanophycin is thus exceedingly rich in nitrogen, having an N:C ratio of about 1:2, which is about an order of magnitude larger than that in typical proteins. While cyanophycin may serve as a carbon-storage compound, its principal role is therefore believed to serve as a nitrogen reservoir.

Under LD conditions, Cyanothece fix nitrogen in the dark, as measured by the nitrogenase activity, and store glycogen during the day [12]. Also in continuous light [12] or continuous dark conditions [13], the nitrogenase activity and cyanophycin storage peak during the subjective night while glycogen storage peaks during the subjective day, indicating the presence of a circadian clock that coordinates these activities. Interestingly, under LD conditions, *Cyanothece* exclusively grows during the day [14], but in continuous dark, when grown on glycerol [13], it still predominantly grows during the subjective day, although Fig. 8 of Ref. [13] leaves open the possibility it may also grow during the subjective night.

These physiological rhythms of *Cyanothece* are mirrored by circadian rhythms in gene expression [15–20]. About 30% of the 5000 genes examined exhibit oscillating expression profiles [15]. Moreover, these genes are primarily involved in core metabolic processes, such as photosynthesis, respiration, energy metabolism, and amino-acid biosynthesis [15]; in contrast, most genes involded in transport, DNA replication and repair, were not differentially expressed [15]. Importantly, genes associated with nitrogen fixation are primarily expressed in the dark, while those underlying photosynthesis are upregulated during the ligth and downregulated during the dark period [15]. Proteomic analysis using partial metabolic heavy isotope labeling identified 721 proteins with changing levels of isotope incorporation [18], of which 425 proteins matched the previously identified cycling transcripts [15]. In particular, the nitrogen fixation proteins were most abundant during the dark [18] while many proteins involved in photo-synthesis are present in higher abundance during the light. Interestingly, proteins involved in storing glycogen, such as the glycogen synthase, peak during the light, while enzymes involved in glycogen metabolism, such as glycogen phosphorylase, GlgP1, have higher levels during the dark [15]. Conversely, the cyanophycin processing enzyme cyanophycinase, CphB, which breaks down cyanophycin into arginine and aspartic acid, shows higher synthesis in the light [18], although, perhaps surprisingly, cyanophycin synthetase, dCphA, appears not to be strongly coupled with the light-dark cycle.

These transcriptomic and proteomic analyses [15–18, 20], together with large-scale computational modeling of the metabolic network [21], provide detailed information about the proteome repartitioning dynamics during the 24 h period. Yet, many questions remain open: First and foremost, why do cyanobacterial cells typically exclusively grow during the day? Cyanobacterial cells have the components to grow at night, which suggests that the strategy to not grow during the dark arises from a cellular trade-off that maximizes the growth rate over 24h [21]. Can this trade-off be quantified, and do cellular growth laws predict that it is optimal to not grow at all during the night? Secondly, in the absence of active protein degradation, the timescale for the relaxation of the proteome is given by the growth rate [4, 5, 22], while at the same time the growth rate of these cyanobacterial cells is affected by how fast the proteome can adjust to the changing light and nutrient levels (glycogen and cyanophycin). This observation is particularly pertinent, because the growth rate of these cyanobacterial cells tend to be low, with cell-division times that are typically longer than 10 hours [6–8]. How much is the growth rate limited by the slow relaxation of the proteome? Thirdly, cyanbacterial cells have a circadian clock, which allows them to predict and anticipate the changes in light and nutrient levels. In general, anticipation becomes potentially beneficial especially when the cellular response is slow [23]. Does anticipation allow cyanobacterial cells to significantly raise their growth rate?

To address these questions, we employ the framework developed by Hwa and coworkers for quantifying the relationship between growth and proteome composition [1–5] and extend it to describe circadian metabolism. This framework is inspired by two key observations: On the one hand the response to a changing environment tends to be extremely complex at the molecular level, involving a myriad of signaling and metabolic pathways. On the other hand, it tends to be global, meaning that in response to a nutrient limitation certain subsets of enzymes are upregulated while others are downregulated. The system is therefore not described in terms of the detailed signaling and metabolic networks, but rather via coarse-grained protein sectors. Each sector contains a subset of enzymes, which share a common purpose, according to the supply-and-demand picture of protein synthesis [3]. Each sector is described by a single coarse-grained enzyme, which can be thought of as representing the average activity of the enzymes in that sector. It is this coarse-grained description that allows for a quantitative mathematical analysis. The framework has been used to describe the effect of protein overexpression [1], cAMP-mediated catabolite repression [2], growth bistability in response to anti-biotics [24], and methionine biosynthesis [25]. And importantly for our analysis, it has recently been extended to describe the transient relaxation dynamics of the proteome in response to a nutrient shift [4, 5]. While these studies have focused on the bacterium *Escherichia coli*, we here employ this framework to study circadian metabolism of cyanobacteria.

The model that we present aims to describe the circadian metabolism of cyanobacteria like *Cyanothece*, which fix carbon during the day and nitrogen during the night, although it can straightforwardly be amended to describe the metabolism of cyanobacteria such as *Synechococcus* and *Synechocystis* that only fix carbon. Arguably the most minimal model to capture the interplay between carbon and nitrogen fixation is one that consists of a ribosome sector, a carbon sector and a nitrogen sector. However, to capture the fact that storing glycogen and cyanophycin does not directly contribute to growth, but only indirectly, by providing carbon and nitrogen the next part of the day, our model also contains two other protein sectors: a glycogen and a cyanophycin synthesis sector. Our model therefore naturally includes two important consequences of building a carbon and nitrogen reservoir: 1) it requires the synthesis of enzymes that do not directly contribute to growth, and hence lower the instantaneous growth rate [1]; 2) storing carbon and nitrogen atoms drains carbon and nitrogen flux away from protein synthesis. Our model further incorporates the dynamics of the carbon and nitrogen reservoirs (glycogen and cyanophycin), the slow relaxation of the proteome in response to the changing nutrient levels, and the capacity to anticipate the changing nutrient levels by mounting a response ahead of time.

We first use this model to study the optimal strategy that maximizes the growth rate over 24 hours. Our analysis reveals that the need to store carbon and nitrogen tends to generate an extreme strategy, in which cells predominantly grow during the day, as observed experimentally [13, 14]. However, our analysis also reveals that the slow relaxaton of the proteome, arising from the slow growth rate, puts a severe constraint on implementing this optimal strategy. In essence, to store enough cyanophcyin during the night to fuel growth during the day, the cyanophycin-storing enzymes need to be expressed at levels that cannot be reached if the cells would only start expressing these enzymes at night. Indeed, to implement the optimal strategy, the cells need to express these enzymes already before the beginning of the night, when they still grow significantly. Interestingly, recent transcriptomics and proteomics data provide evidence for this prediction [20].

## THEORY

The central ingredients of the framework of Hwa and coworkers [1–5] are the coarse-grained protein sectors and the balance of fluxes between them. We describe these elements in turn.

**Protein setors** The sectors are defined experimentally by how the enzyme expression levels vary with the growth rate in response to different types of nutrient limitation [3]. The C-sector is defined as the subset of enzymes whose expression levels increase as the growth rate decreases upon a Carbon limitation, yet decrease as the growth rate decreases upon a nitrogen limitation or translation inhibition [3]. A mass-spec analysis of *E. coli* revealed that this sector contains enzymes involved in ion-transport, the TCA- cycle and locomotion [3]. The A-sector is defined as the group of proteins that are upregulated in response an A-limitation—a nitrogen limitation— yet downregulated in response to carbon or translation limitation. In *E. coli*, this sector consists of enzymes that are involved in the incorporation of nitrogen into amino-acids [3]. The R-sector contains the ribosomes, which increase in abundance as the growth rate decreases upon the addition of a translation inhibitor. The study of Hui and coworkers on *E. coli* also identified an S-sector, consisting of enzymes whose expression levels increase in response to both carbon and nitrogen limitation, and a U-sector, consisting of proteins that are uninduced under any of the applied limitations [3].

In our model, we are interested in the interplay between carbon and nitrogen assimilation, and the simplest model that can capture this interplay is one that considers an R-sector, a C-sector and an A-sector. The mass fractions of the proteins in these sectors are denoted by *ϕ*_R_, *ϕ*_C_ and *ϕ*_A_, respectively. Our model does not explicly contain an S- and a U-sector, although we emphasize that as experimental data becomes available the model can straightfowrardly be extended to include these sectors [3]. Following Hui *et al*., we also stress that these sectors are ultimately defined experimentally [3]. In our case, the C-sector is defined as consisting of those enzymes that are upregulated in response to a carbon limitation, yet downregulated in response to an A- or R-limitation. The carbon limitation can be in the form of reduced CO_2_ and light levels during the day, but also reduced glycogen levels during the night. Our model thus lumps all proteins that are involved in providing carbon skeletons for amino-acid synthesis into one sector, the C-sector. We anticipate that this sector contains enzymes of not only the photosynthesis machinery, but also the TCA cycle, as well as enzymes involved in degrading glycogen, such as glycogen phosphorylase GlgP. Experiments need to establish whether it would be necessary to split this C-sector up into separate sectors for, e.g., photosynthesis, glycogen breakdown and downstream carbon processing (e.g. TCA cycle).

Similarly, we define the A-sector as the set of enzymes that are upregulated in response to nitrogen limitation, yet downregulated in resopnse to a C- or R-limitation. We envision that nitrogen limitation can be imposed by reducing N_2_ levels, by employing a titratable nitrogen uptake system [3], or by lowering levels of cyanophycin. While, again, experiments need to identify which enzymes belong to this sector, we expect that it contains not only the nitrogenase enzymes that reduce N_2_ into ammonia and the enzymes that subsequently incorporate the nitrogen into amino-acids, but also the proteins that are involved in the breakdown of cyanophycin, such as cyanophycinase CphB.

Following Scott *et al*., the model also includes an unresponse fraction *ϕ*_Q_, although this parameter will be absorbed in the maximal ribosomal fraction *ϕ*_R,max_, as described below [1–3].

**Storing fractions** While this model is indeed highly coarse-grained, we do consider two other sectors, which contain enzymes that store glycogen during the day and nitrogen during the night. Their fractions are denoted by *ϕ*_SC_ and *ϕ*_SA_, respectively. Enzymes belonging to *ϕ*_SC_ are glycogenin and glycogen synthase, which are indeed upregulated during the day [17, 18]. Cyanophycin is synthesized from arginine and aspartate by a single enzyme, cyanophyinc synthetase, CphA, which thus forms the *ϕ*_SA_ sector [17]. A key point is that expressing the glycogen-storing enzymes slows down growth during the day yet enables growth during the night, while expressing cyanophycin synthetase sloww down growth during the night, yet enables growth during the day. Even though the growth laws are linear, this creates a feedback between growth at night and during the day that yields a non-linear response, as we discuss in more detail below.

**Steady-state flux balance** The experiments by Hwa and coworkers on *E. coli* have revealed that the steady-state growth rate varies linearly with the size of the protein sectors [1–3]. These linear relationships can be understood by combining the following ideas: a) in steady-state the fluxes *J_α_* through the different sectors α= R, C, A are balanced, so that there is no build-up of intermediates like amino-acids; b) the growth rate *λ* is proportional to the flux through the sectors; c) the flux through a sector scales linearly with the size of the sector. Combining these ideas makes it possible to quantitatively describe the experiments of [2, 3], explaining how each sector is upregulated in response to one type of limitation, while downregulated in response to another type of limitation [2, 3]. Moreover, the model can quantitatively describe how the growth rate decreases as an unnecessary protein, which does not directly contribute to growth, is expressed via an artificial inducer [1]. The latter is important, because the proteins that store glycogen and cyanophycin, respectively, can be thought of as proteins that do not directly contribute to growth; they only contribute by providing the carbon- and nitrogen sources for the next part of the day. Our model incorporates these three ingredients a)-c), but adds a fourth, d): a certain fraction of the flux through the carbon and nitrogen sector is reserved for storing glycogen during the day and cyanophycin during the night, respectively.

While our full model is time dependent, we will first consider a simpler model in which we can directly use the growth laws derived by Hwa and coworkers [1–3]. Specifically, during the day the principal source of carbon is CO_2_, while that of nitrogen is cyanophycin, which decreases with time. During the night, the principal source of nitrogen is N_2_, while that of carbon is glycogen, which falls with time. As a result, the growth rate *λ* will, in general, be time-dependent, *λ* = *λ*(*t*). Moreover, because the glycogen and cyanophycin concentrations vary with time, the proteome fractions, which are adjusted in accordance with the nutrient availability, will change not only upon the shift from day to night, but also continue to change throughout the day and night. As was pointed out in [22] and also in [4, 5], in the absence of active protein degradation, the proteome relaxes with a timescale that is set by the growth rate *λ*(*t*). This means that when the growth rate is low, the proteome will relax slowly, and may not be in quasi-equilibrium with respect to the instantaneous levels of glycogen during the night and cyanophycin during the day. Below we will take this slow relaxation of the proteome into account. However, to introduce the main elements of the model, it will be instructive to first assume that the growth rate is so high, that the proteome is always in quasi-equilbrium with respect to the instantaneous nutrient levels, set by the CO_2_/light and cyanophycin levels during the day, and the glycogen and N_2_ levels during the night. The growth thus depends on time, but not explicitly, and only implicitly via the levels of glycogen and cyanophycin: *λ*(*t*) = *λ*([C](*t*), [N](*t*)), where [C](*t*) and [N](*t*) are the time-dependent carbon and nitrogen sources. We call this model the quasi-equilibrium model.

**Quasi-equilibrium model** The first two ingredientsa) and b) imply that in the quasi-equilibrium model

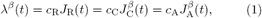

where *c_α_* are the stochiometric requirements for cell growth [3]. Here, we have added the superscript *β* = L, D, with L standing for light and D for dark, to remind ourselves that the fluxes through the carbon and nitrogen sector and thereby the growth rate, depend on the source of carbon and nitrogen used, which differs between day and night.

The third observation c) means that for the ribosomal sector

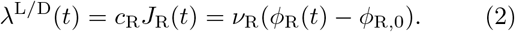

Here, *v*_R_ = *c*_R_*k*_R_, where *k*_R_ describes the translation efficiency, which can be varied experimentally using a translation inhibitor such as cloramphenicol [1–3]. The quantity *ϕ*_R,0_ is the fraction of ribosomes that is not active in steady-state, yet can become active during the transition from one environment to the next [4, 5]. In the quasi-equilibrium model considered here, it is a constant, independent of time.

The ingredients c) and d) imply that the flux through the carbon sector that flows into the other sectors is given by

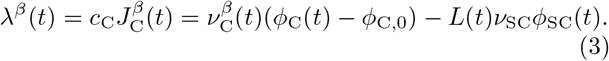

Here, *L*(*t*) is an indicator function that is 1 during the day and 0 during the night. Indeed, during the day both terms are present. The first term on the right-hand side describes the carbon flux that would flow into the other sectors if no carbon were stored into glycogen during the day. The second term indeed describes the flux that is not used for growth during the day, but rather lost in storing glycogen. During the night, no glycogen is stored and the second term is absent. In the first term,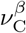 is a measure for the efficiency of the carbon sector. It depends on the quality and the amount of nutrient [2, 3], but can also be varied experimentally—in *E. coli* by titrating a key enzyme such as the lactose permease [2, 3]. In our model,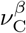depends on the part of the day, as indicated by *β* = L,D: during the day, the principal carbon source is CO_2_, which means that the value of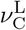 will depend on the concentration of CO_2_ and light levels. Since we will model the light intensity as a step function, during the day the light level and hence 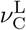(*t*) is constant, and equal to 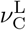(*t*) = 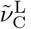. In contrast, during the night, the principal source of carbon is glycogen, which decreases during the night. This affects the carbon-processing efficiency. We will model this as

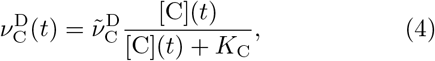

where [C](*t*) is the time-dependent concentration of glycogen and *K*_C_ is the glycogen concetration at which the enzyme efficiency is reduced by a factor of 2. The quantiy 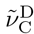is the maximal efficiency of the carbon-sector with glycogen as the carbon source; it does not depend on time. The quantity *ϕ*_C,0_ is the fraction of carbon-processing enzymes that is not used for growth. For *E. coli* it is very close to zero, and from hereon we assume it to be zero. The quantity *v*_SC_ describes the efficiency of the glycogen-storing enzymes, and is taken to be constant.

For the nitrogen-sector, we similarly obtain

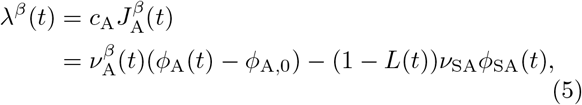

where in the calculations performed here we assume that *ϕ*_A,0_ is zero, even though the experiments indicate that for *E. coli* the unused fraction in the A-sector is about 10% [3]. The nitrogen-processing efficiency during the day depends on the concentration of stored cyanophycin, [N], via

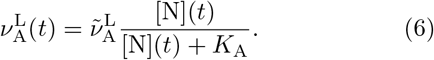

The nitrogen-uptake efficiency during the night depends on the amount of N_2_, which we assume to be constant throughout the night. The efficiency is thus given by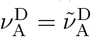

Combining all four ingredients a) – d), i.e. Eqs.1–6, yields

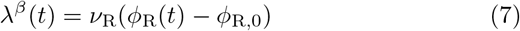

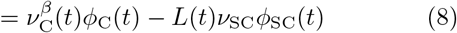

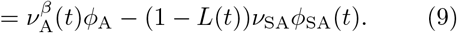

**Proteome balance** The protein sectors obey at all times t the constraint

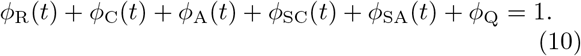

The growth rate λ will be maximal, λ → λ_max_, when the storing, carbon- and nitrogen-processing fractions approach zero, and the ribosomal fraction becomes maximal

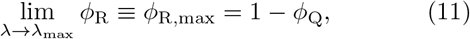

allowing us to rewrite the constraint as:

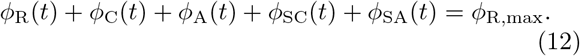

We note that this definition of *ϕ*_R,max_ differs slightly from *ϕ*_max_ defined in Ref. [2, 3].

**Growth rate in quasi-equilibrium model** In our model, the input parameters are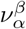 and *ϕ*_R,0_, while the storing fractions *ϕ*_SC_(*t*) and *ϕ*_SA_(*t*) are control parameters over which we will optimize to maximize the growth rate over a 24h period. In the quasi-equilibrium model, the optimal *ϕ*_SC_ during the night is zero and the optimal *ϕ*_SA_ during the day is zero. In this model, we thus have one optimization parameter *ϕ*_SC_ for the day, and another for the night, *ϕ*_SA_. In this quasi-equilibrium model, the other protein sectors relax instantaneously, to values that, for the day, are determined by the efficiencies *v*_R_, 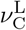, the instantaneous efficieny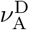(*t*)and the optimization parameter *ϕ*_SC_(*t*), and, for the night to values given by*v* _R_,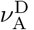, the instantaneous value of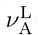(*t*) and the optimization parameter *ϕ*_SA_(*t*). The 4 equations, Eqs. 7–9 together with the constraint Eq. 12, thus contains 4 unknowns *ϕ*_R_*, ˚*_C_*, ˚*_A_,λ, which can be solved to obtain the instantaneous growth rate for the day and night, respectively, for the quasi-equilibrium model:

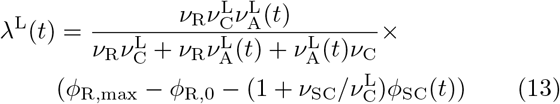

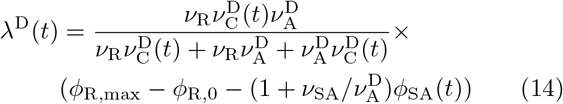

Clearly, during the day the growth rate, for given CO_2_ and cyanophycin levels, is maximal when no glycogen is stored and *ϕ*_SC_ = 0. This defines a maximum growth rate during the day

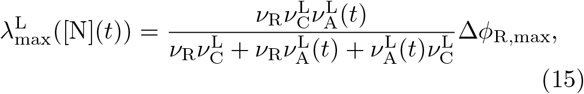

where Δ*ϕ*_R,max_ =*ϕ*_R,max_ − *ϕ*_R,0_. The maximal growth rate depends on the instantaneous amount of cyanophycin, [N](*t*), because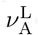(*t*)depends on [N](*t*) (see Eq. 6). From Eq. 13 we find the storing fraction 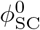that reduces the growth rate to zero during the day:

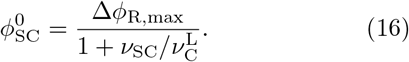

This allows us to rewrite Eq. 13 as

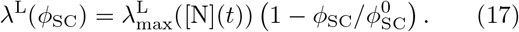

Equivalently, we find for the growth rate during the night

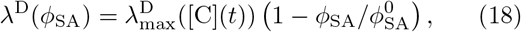

with 

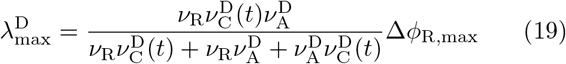

and 

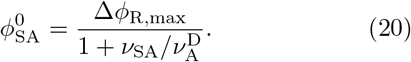

A few points are worthy of note. Firstly, Eqs. 17 and 18 show that the growth rate decreases linearly with*ϕ*_SC_ and *ϕ*_SA_, respectively. In fact, Scott *et al*. derived a similar relation for the growth rate when an unnecessary protein is expressed [1]. This highlights the idea that storing glycogen and cyanophycin reduces the growth rate, because synthesizing these storage molecules requires proteins that do not directly contribute to growth—thus taking up resources that could have been devoted to making more ribosomes. Indeed, building carbon and nitrogen reservoirs only pays off the next part of the day, which can be seen by noting that the maximum growth rate during the day, 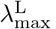, increases with the amount of cyanophycin that has been stored the night before (via 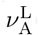, see Eq. 6), while the maximum growth rate during the night, 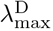 increases with the amount of glycogen that has been stored the day before (via 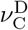, see Eq. 4). Clearly, the cell needs to strike a balance between maximizing the instantaneous growth rate and storing enough resources to fuel growth the next part of the day.

However, there is also another effect: building the reservoirs reduces the growth rate not only because it requires proteins that do not directly contribute to growth, but also because it drains carbon and nitrogen flux. This manifests itself in the intercepts 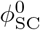 and 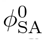 at which the growth rate is zero (see Eq. 16). This effect puts a hard fudnamental bound on the maximum rate at which glycogen and cyanophycin can be stored. For glycogen the maximum storing rate is given by

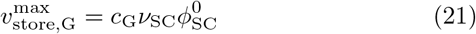

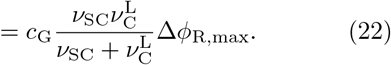

Here, *c*_G_ is a stochiometric coefficient that reflects the number of carbon atoms that are stored in a glycogen molecule. This expression shows that the maximal storing rate 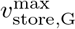 increases with Δ*ϕ*_R,max_. This is because Δ*ϕ*_R,max_ limits the fraction of the proteome that can be allocated to storing glycogen, 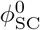. The expression also reveals that 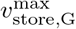 depends on*v* _SC_ and 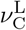. The maximum storing rate 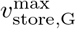 initially increases with*v* _SC_, simply because that increases the rate at which the glycogen storing enzymes operate. However, the increased flux of carbon into glycogen also means that less carbon is available for making the glycogen-storing enzymes themselves. As a result, as*v* _SC_ increases, the maximal fraction 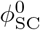 of glycogen-storing enzymes decreases (see Eq. 16). In the limit that*v* _SC_ becomes very large, i.e. much larger than*v*_C_, then 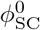 becomes zero, and the rate at which glycogen is stored becomes independent of *v*_SC_. In this regime, all the carbon flows into glycogen and the storing rate instead becomes limited by 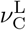, 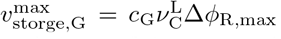. In this limit, *ϕ*_R_ = *ϕ*_A_ = 0 and *ϕ*_C_ = Δ, such that there is no carbon flow devoted to growth, 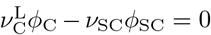 (Eq. 3), but only to storing glycogen. As we will see below, this will put a strong constraint on the maximal growth rate of the cyanobacteria.

**Reservoir dynamics** The growth rate depends on the efficiencies 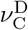(*t*) and 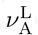(*t*), which depend on the amount of glycogen and cyanophycin, respectively (see Eqs. 4 and 6). The dynamics of their concentration is given by

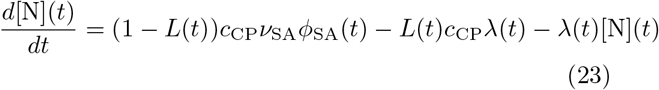

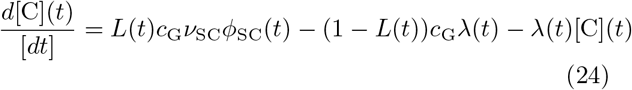

The last term in both equations is a dilution term, where we have exploited that cells grow exponentially with rate λ(*t*). The first term describes the accumulation of the stores due to the storing enzymes, with *c*_CP_*, c*_G_ being stoichiometric coefficients that reflect how many nitrogen and carbon atoms are stored in a cyanophycin and glycogen molecule, respectively. The second term describes the consumption of cyanophycin and glycogen that fuels growth. Focusing on glycogen, this term can be understood by noting that the depletion of glycogen during the night is given by the rate at which the carbon sector consumes glycogen:

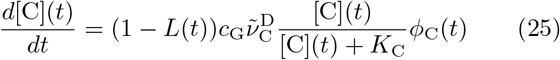

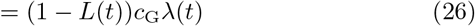

where in the second line we have exploited that in quasi-equilibrium the growth rate λ (*t*) is given by the flux through the carbon sector (see Eqs. 3 and 4). Importantly, this expression reveals that the depletion of the store depends on the growth rate not only because that sets the dilution rate (reflected by the third term in Eqs. 23 and 24), but also because the growth rate sets the rate at which the store is consumed (second term).

**Slow proteome dynamics** The proteome will in general not be in quasi-equilibrium with respect to the instantaneous nutrient levels. To take into account the relaxation of the proteome we first define the mass fractions *ϕ_α_* of the different sectors

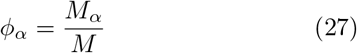

where *M_α_* is the protein mass of sector *α* = R, C, A, SC, SA, Q and *M* is the total mass. The total rate at which proteins are synthesized is given by 

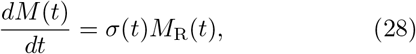

where *M*_R_ is the mass of the ribosomal sector, consisting of the mass of the ribosomes and the ribosome-affiliated proteins [5]. The quantity *σ*(*t*) is the instantaneous translational efficiency. It corresponds to the average translational efficiency, and does not distinguish between active and inactive ribosomes [5]. When we divide the above equation by *M(t)*, we obtain the instantaneous growth rate [5, 22]

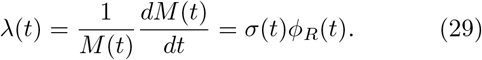

To obtain the evolution of the different protein sectors, we denote the fraction of the number of ribosomes that are allocated to making protein sector *α* as *χ_α_*. The evolution of the proteome mass *M_α_* is then 

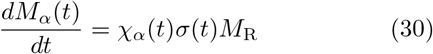

and that of the proteome fraction [5, 22] 

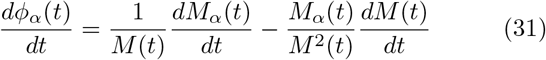

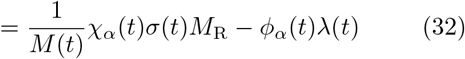

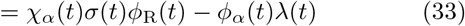

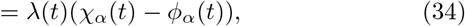

where in going to the last line we have exploited Eq. 29. This equations shows that when *χ_α_* (*t*) adjust rapidly to a new nutrient environment, as recent experiments indicate [4, 5], the relaxation of the proteome is dominated by the growth rate λ (*t*). Importantly, the equation also shows that when *χ_α_* (*t*) = *ϕ_α_* (*t*), the proteome has equilibrated: the fractions no longer change with time.

Recent experiments indicate that after a nutrient up-shift the translational efficiency *σ*(*t*) and the fraction *χ_α_* (*t*) of ribosomes devoted to making proteins of sector*α* rapidly approach their new steady-state values as set by the new environment [4, 5]. We therefore make the simplification [22], also used in [5], that after a day-night (and night-day) transition *σ*(*t*) immediately takes the final value *σ\** set by the new environment and that *χ_α_* (*t*) immediately takes the value of the final fraction 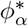 in the new environment. However, in our system, the amounts of glycogen and cyanophycin change with time, and the proteome fractions continually adjust to this. The “final” fractions 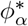 are thus target fractions that themselves change with time, and similarly for the translation efficiency *σ*:

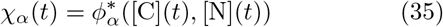

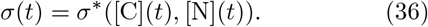

These quantities are set such that if *ϕ_α_* (*t*) were equal to 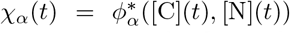 and *σ*(*t*) were equal to *σ\** ([C](*t*), [N](*t*)), the fluxes through the different sectors would be balanced and the growth rate would be equal toλ* (*t*):

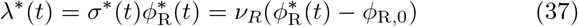

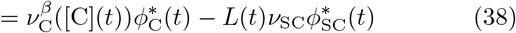

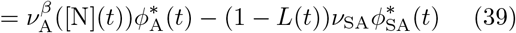

Importantly, we do not only need to consider the target fractions for the R-, C-, A-, and Q-sector, but also for the storing fractions:χSC(*t*) = 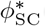(*t*) and χSA(*t*)= 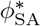(*t*). Eqs. 37–39 are thus solved subject to the following constraint

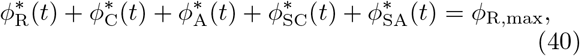

where *ϕ*_SC_(*t*) and *ϕ*_SA_(*t*) are optimization parameters described in more detail below. This equation states that the total ribosome protein synthesis fraction 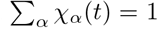 at all times, which guarantees that 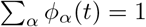 at all times.

**Anticipation** The cells need to repartition its proteome every 12h as the cells move from day to night, and vice versa. Moreover, the cells need to continually adjust its proteome to the changing levels of cyanophycin and glycogen. However, the relaxation of the proteome is, in the absence of protein degradation, set by the growth rate (see Eq. 34), which for cyanobacteria, with cell division times in the range of 10 – 70h, is low compared to the 24 hr period of the day-night cycle. This slow relaxation of the proteome will tend to make the growth rate suboptimal. Interestingly, cyanobacteria, ranging from *Synechococcus*, *Synechocystis*, to *Cyanothece* have a circadian clock, which allows them to anticipate the changes between day and night and to adjust their proteome ahead of time.

To include this into the model, we introduce the notion of the anticipation time *T*_a_. That is, the cells will compute the target protein fraction 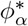(*t*) at time *t* (see Eqs. 37–39) based on the values of 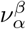 (*t* + *T*_a_) at the later time *t* + *T*_a_. The ribosome fraction *χ_α_* (*t*) = 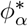(*t*) devoted to making proteins of sector*α* at time *t* is thus determined by the protein efficiencies 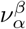 (*t* + *T*_a_) at a later time *t* + *T*_a_. This allows cells to already adjust their proteome before the end of the day (night) is over, and steer it towards the target protein fractions set by the efficiencies 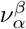(*t* + *T*) in the following night (day).

There is one subtlety, which we address in a rather adhoc fashion. The efficiencies 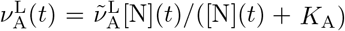 and 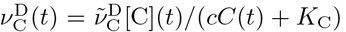depend on the concentrations of cyanophycin and glycogen at time *t*, respectively. Experiments on plant cells in combination with modeling [26] suggest that cells might be able to extrapolate the current concentrations [C](*t*) and [N](*t*) to estimate the concentrations at time *t*+*T*, [C](*t*+*T*) and [N](*t*+*T*), respectively. While this could be included into our model, we make the simplication that the cells base the future efficiency based on the current concentration of the store.

The target fractions 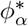(*t*) are thus obtained by solving Eqs. 37–39 but with the protein effciencies given by

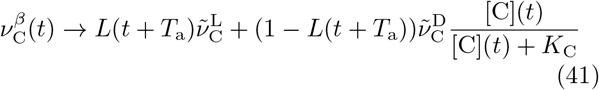

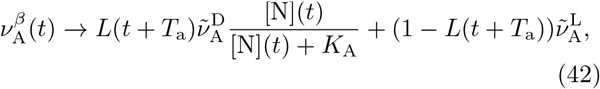

where, as before, *L*(*t*) is an indicator function that is 1 during the day and 0 during the night.

In addition, in this anticipation model, we also take into account that the protein storing fractions *ϕ*_SC_ and *ϕ*_SA_ can be made ahead of time: the synthesis of the glycogen-storing enzymes can already start before the beginning of the day, while the production of the cyanophycin-storing enzyme can already start before the beginning of the night. As we will see, especially the latter can significantly enhance the growth rate. Importantly, while the storing enzymes are synthesized ahead of time, we assume that they become active only when they need to be, i.e. the cyanophycin-storing enzymes are active only during the night, while those storing glycogen are only active during the day.

**Overview full model** The model that takes into the slow proteome relaxation dynamics but *not* anticipation, is given by Eq. 29 which gives the instantaneous growth rateλ(*t*), Eq. 34 that describes the evolution of *ϕ_α_* (*t*), and Eqs. 35–40, which are solved to yield *χ_α_* (*t*) = 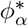(*t*) in Eq. 34 and *σ*(*t*) = *σ\** (*t*) in Eq. 29, together with the dynamics for the concentrations of cyanophycin and glycogen, Eqs. 23 and 24. Moreover, in this so-called slow-proteome model, we set *ϕ*_SC_ to be zero during the night and *ϕ*_SA_ to be zero during the day, and optimize over the magnitude of their values during the day and night, respectively.

The full model, called the anticipation model, is based on the idea that the cell possesses a clock that not only makes it possible to anticipate the changes in protein efficiencies 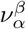(*t*) between day and night, but also to express the protein storing fractions in an anticipatory fashion. The full model is thus exactly the same as the slow-proteome model, except for the following two ingredients:1) the efficiencies 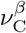(*t*) and 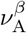(*t*) in Eqs. 37–38 are replaced by those of Eqs. 41 and 42; 2) the protein-storing fractions *ϕ*_SC_(*t*) and *ϕ*_SA_(*t*) are optimized not only with respect to their magnitude, but also with respect to the timing of their expression.

## PARAMETER SETTINGS

In our model, the key parameters that can be varied experimentally are 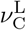, which is determined by the CO_2_ and light levels during the day, 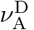, which is by the N_2_ level, and *v*_R_, which can be varied experimentally via a translational inhibitor such as chloramphenicol. The parameters
 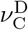and 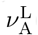 are set by the nutrient quality of glycogen and cyanophycin, respectively, while*v* _SC_ and*v* _SA_ are determined by the efficiencies of the glycogen and cyanophycin storing enzymes, respectively. We will keep these parameters constant in all the results that we present below. The parameters are set such that for the baseline parameter values the average cell-division time is roughly 24h. The values of *ϕ*_R,max_ and *ϕ*_R,0_ are inspired by those measured for *E. coli* [3]. The parameters *ϕ*_SC_ and *ϕ*_SA_ are optimization parameters, as described above. We optimize these parameters by numerically propagating our model for given values of *ϕ*_SC_ and *ϕ*_SA_ and numerically computing the average growth rate 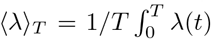 over one period of duration *T*, which under normal conditions is *T* = 24h.

## RESULTS

### Quasi-equilibrium model

It is instructive to first consider the scenario in which the relaxation of the proteome is instantaneous, such that at any moment in time the protein fractions are optimally balanced based on the values of the protein efficiencies 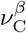 and 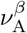 and the instantanlevels of glycogen, [C](*t*), and cyanophycin, [N](*t*), repspectively.

Fig. 1A shows a heat map of the average growth rate over 24h, 〈λ〉_24_ as a function of the fractions of proteins that store glycogen and cyanophycin, *ϕ*_SC_ and *ϕ*_SA_, respectively. The parameters have been set such that the system is symmetric,*v* _SC_ =*v* _SA_, 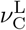 = 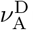, *K*_C_ = *K*_A_, *c*_G_ = *c*_CP_, except that the maximum growth rate during the day is slightly larger than that during the night because 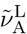 = 6/h is slightly larger than 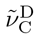 = 2/h. The prominent feature of the figure is that even though the system is slightly asymmetric, meaning that the system could grow during the dark, the storing fractions that maximize the growth rate are such that the optimal cyanophycin-storing protein fraction 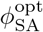 is markedly non-zero, while the optimal glycogen-storing protein fraction, 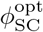, is essentially zero.

**FIG. 1:**
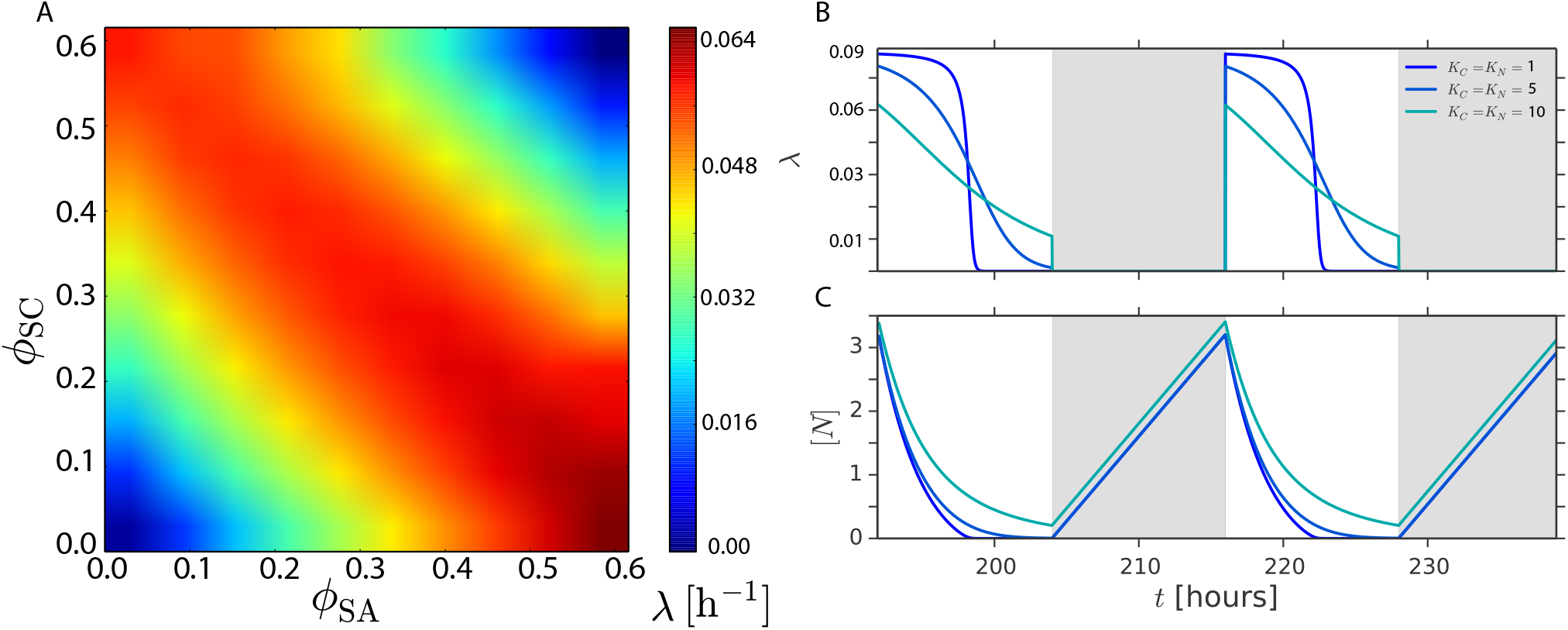
Optimal growth strategy in the quasi-equilibrium model. In this model, the proteome fractions relax instantaneously and as such are always in quasi-equilibrium with the instantaneous levels of glycogen, [C](*t*), and cyanophycin, [N](*t*). In this model, the instantaneous growth rate is given by Eqs. 13 and 14 (or equivalently Eqs. 17 and 18), while the reservoir dynamics is given by Eqs. 23 and 24. The model is nearly symmetric between day and night, 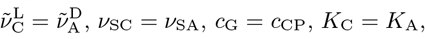 except that 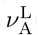 = 3/h is slightly larger than 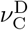 = 1/h. (A) Heat map of the average growth rate over 24h, 〈*λ*〉_24_, as a function of *ϕ*_SC_ and *ϕ*_SA_. The heatmap is obtained by numerically propagating Eqs. 23 and 24, with λ(*t*) given by Eqs. 13 and 14, for different values of *ϕ*_SC_ and *ϕ*_SA_; the average growth rate is obtained by numerically evaluating 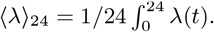 It is seen that there exists a combination of storing fractions that maximizes the growth rate, 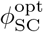 and 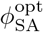; moreover, 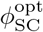 is close to zero, while 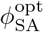 is close to the maximal fraction 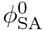 at which the growth rate becomes zero, see Eq. 20. (B) Time traces of λ(*t*) at 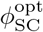 and 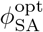, not only for *K*_C_ = 5*c*_G_ = *K*_A_ = 5*c*_CP_, as in panel A, but also for two other values. Clearly, the cells only grow during the day. The growth rate during the night is zero, because the storing fraction 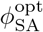 is close to the maximal fraction 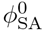 at which the growth rate is zero. This shows that the the optimal strategy in the quasi-equilibrium model is to store as much cyanophycin as possible during the night, because that maximizes the growth rate during the day. The explanation of this behavior is given in Fig. 2. (C) Time traces of the cyanophycin levels for different values of *K*_C_ = *K*_A_. Parameter values: 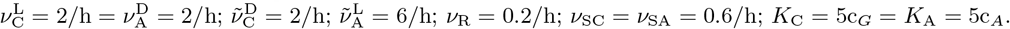

To elucidate Fig. 1A, we show in panel B the growth rate*λ* (*t*) of the system with the optimal storing fractions *ϕ*^opt^_SC_ 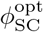 and 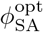, for three different values of *K*_C_ = *K*_A_. Strikingly, the growth rate is zero during the night. The cells only grow during the day, even though with these parameters the cells would have the capacity to grow during the night, had they not to store so much cyanophycin. Indeed, the optimal storing fraction 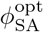 that maximizes the growth rate is close to the fraction 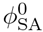 at which the growth rate becomes zero, see Eq. 20.

The mechanism that underlies the optimal strategy is illustrated in Fig. 2. Panel A shows the average growth rate during the day*〈λ 〉*_L_ as a function of *ϕ*_SC_ for different values of *ϕ*_SA_, while panel B of this figure shows the average growth rate during the night,*〈λ 〉*_D_, as a function of *ϕ*_SA_ for different values of *ϕ*_SC_. First of all, note that the maximum growth rate during the day is only slightly larger than that during the night—the asymmetry between day and night is indeed (chosen to be) weak. Yet, the optimal strategy, which maximizes the average growth rate over 24h, is to not grow at all during the night. To understand this, note that storing more cyanophycin during the night will enhance the growth rate during the day (panel A), yet lower it during the night (panel B). Similarly, storing more glycogen during the day will raise the growth rate during the night (right B), yet lower it during the day (panel A). The crux is that the cost of storing less glycogen during the day—a lower growth rate at night—decreases when more cyanophycin is stored, while at the same time the benefit of storing more cyanophycin—growing faster during the day—is largest when the amount of stored glycogen is minimal. This tends to favor a strategy where the maximum amount of cyanophycin is stored during the night, while a minimal amount of glycogen is stored during the day. Naturally, the argument also works in the converse direction, yielding a strategy where the maximal amount of glycogen is stored during the day and the minimal amount of cyanophycin is stored during the night. Yet, because the maximal growth rate during the day is larger than the maximal growth rate during the night, the former strategy is favored.

**FIG. 2:**
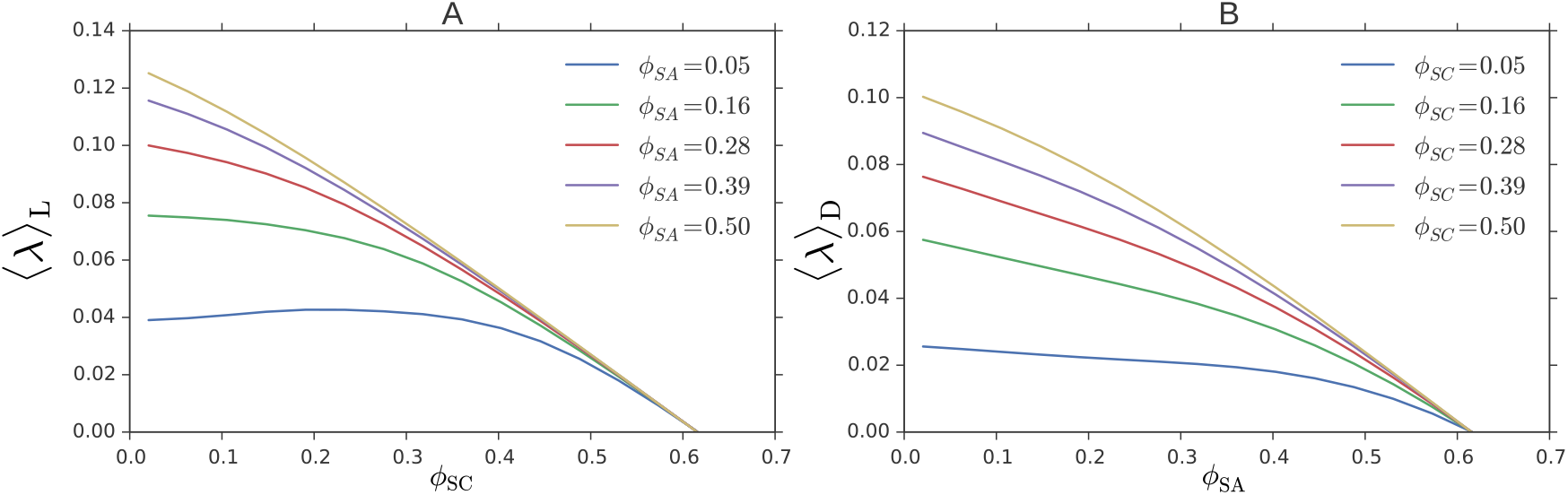
Mechanism underlying the optimal strategy that maximizes the growth rate in the quasi-equilibrium model, given by Eqs. 13 and 14 and Eqs. 23 and 24. (A) The average growth rate during the day, *〈λ〉_L_* as a function of *ϕ*_SC_ for different values of *ϕ*_SA_. (B) The average growth rate during the night, *〈λ〉_D_* as a function of *ϕ*_SA_ for different values of *ϕ*_SC_. These figures have been obtained by numerically propagating Eqs. 13 and 14 and Eqs. 23 and 24 for different combinations of *ϕ*_SC_ and *ϕ*_SA_. The key point is that the cost of storing less glycogen—a lower growth rate at night—decreases when more cyanophycin is stored (and vanishes in fact when *ϕ*_SA_ approaches its maximum 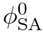 where the growth rate becomes zero), while the benefit of storing more cyanophycin—a higher growth rate during the day—increases as less glycogen is stored during the day (because *ϕ*_SC_ is smaller). This yields an optimal strategy that maximizes the growth rate in which the cells exclusively grow during the day. Parameter values as in Fig. 1: 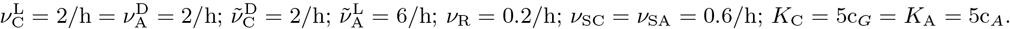Growth rates are in units of 1/h

An important question is how generic this tipping-point strategy in which cells predominantly grow in one phase of the day, is. What is essential is that the maximum growth rate during the day, 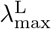 (Eq. 15), is larger than that during the day, 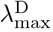 (Eq. 19). Yet, the precise values of the efficiencies 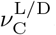, 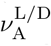 tend to be less important, depending on the values of*v* _SC_ and*v* _SA_. If the model is fully symmetric,*v* _SC_ =*v* _SA_, 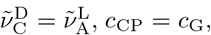except that 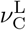 > 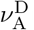, then a tipping-point strategy is still favored, provided that*v* _SC_ and*v* _SA_ are not too large with respect to 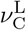 and 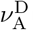, respectively. The reason is that while increasing 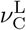 with respect to 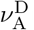 increases the maximum growth rate during the day, which tends to favor growing exlucisely during the day, it also enhances the capacity to store glycogen (as compared to that of storing cyanophycin), which tends to favor growing at night. This effect is particularly pronounced when*v* _SC_ and*v* _SA_ are large compared to 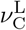 and 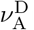, respectively, because then the storing rates become limited by 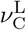 and 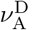, rather than being determined by*v* _SC_ and*v* _SA_, respectively (see discussion below Eq. 22).

The panels of Fig. 2 also reveal that the growth rate is initially fairly constant, before it markedly drops to zero when the storing fraction becomes equal to the maximal storing fraction, given by Eq. 16 for *ϕ*_SC_ (panel A) and Eq. 20 for *ϕ*_SA_ (panel B). The curves deviate from the linear relationship betweenλ and the expression of an unused protein, as found in Scott *et al*. [1]. In fact, also Eqs. 13 and 14 would predict a linear relationship between〈λ〉_L/D_ and *ϕ*_SC_*/ϕ*_SA_ if 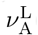(*t*) and 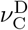(*t*) were constant in time. However, 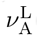 and 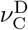(*t*) are not constant in time, because the cyanophycin and glycogen concentrations decrease with time, as can be seen for the cyanophycin concentration in panel C of Fig. 1 (where *ϕ*_SC_ and hence [C] are very small). As a result of this reservoir depletion, also the growth rate varies in time. Moreover, the reservor depletion also underlies the observation that the rise in the growth rate upon decreasing *ϕ*_SC_*/ϕ*_SA_ becomes less pronounced for low *ϕ*_SC_*/ϕ*_SA_ (Fig. 2): in this regime, the growth rate during the day (night) is limited by the amount of cyanophycin (glycogen) during the night (day); decreasing *ϕ*_SC_ (*ϕ*_SA_) only means that the reservoir is depleted more rapidly, yielding no significant net increase in〈λ〉_L_ and〈λ〉_D_; indeed, only by storing more can the growth rate be enhanced further.

The central prediction of this quasi-equilibrium model is thus that the cells do not tend to grow at night, as observed experimentally for *Cyanothece* [14], because that allows it to grow so much faster during the day that the average growth rate over 24h increases. However, this quasi-equilibrium model is based on the assumption that the proteome relaxes instantly, while the relaxation rate, in the absence of protein degradation, is set by the growth rate, which, with typical cell-division times of 10–70h [6–8], is fairly low for cyanobacteria. In fact, to grow faster, the cell needs to store more, while the maximum storing capacity is limited by *ϕ* 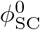 and 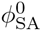, which depend not only on Δ*ϕ*_R,max_, but also on 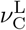 and 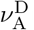, respectively, as discussed below Eqs. 22 and 22. How severe this constraint can be, is seen in panel B of Fig. 1: for the lowest value of *K*_C_ = *K*_A_ shown, the cells grows faster during the beginning of the day. However, because the cyanophycin stored is then depleted more radpily (see panel C below), the growth rate drops sharply well before the end of the day. Here, more cyanophycin can not be stored, simply because *ϕ*_SA_ has already reached its maximum, 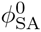. The limited capacity to store thus puts a severe constraint on the growth rate, which limits the proteome relaxation rate. Can the cell under these conditions implement the optimal strategy to maximize the growth rate, as shown in Fig. 1 and Fig. 2? To address this question, we will turn in the next section to the influence of the slow proteome dynamics.

### Slow-proteome model

Fig. 3 shows time traces of the growth rate, the protein fractions and the glycogen and cyanophycin levels for our slow-proteome model. The parameters 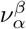 are identical to those of the quasi-equilibrium model corresponding to Fig. 1, yet the magnitudes of *ϕ*_SC_ and *ϕ*_SA_ have been optimized to maximize the growth rate *〈λ〉*_24_ over 24h (the time windows of *ϕ*_SC_ and *ϕ*_SA_ expression have not been optimized in this model, in contrast to in the anticipation model studied in the next section).

**FIG. 3:**
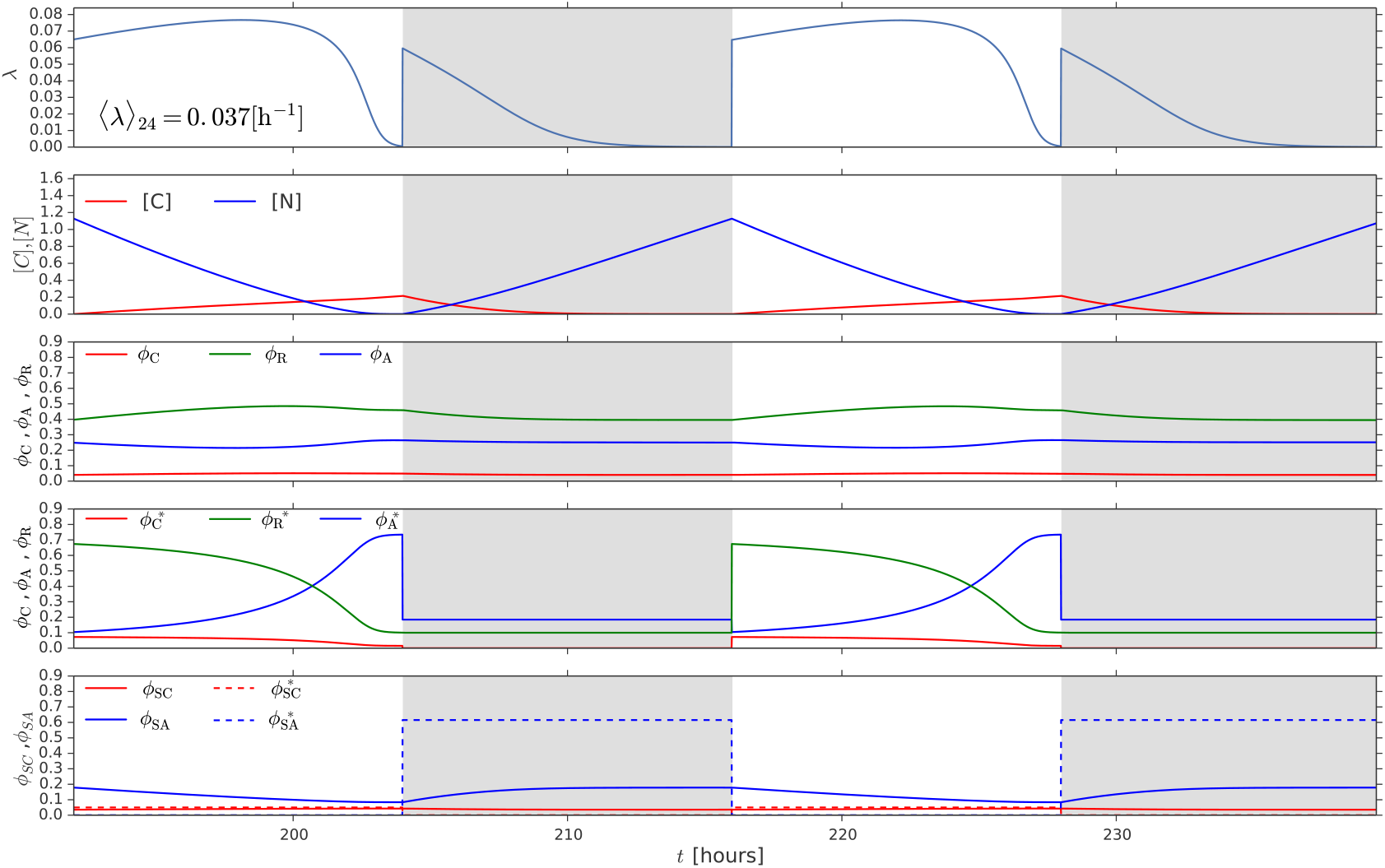
Dynamics of the slow-proteome model, given by Eq. 29 forλ (*t*), Eq. 34 for *ϕ_α_* (*t*), and Eqs. 35–40, which are solved to yield 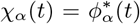 in Eq. 34 and *σ*(*t*) = *σ \** (*t*) in Eq. 29, together with Eqs. 23 and 24 for the reservoir dynamics. Time traces of the growth rate λ (*t*) (1/h, top row), glycogen levels [C](*t*) and cyanophycin levels [N](*t*) (second row), protein fractions *ϕ*_R_(*t*)*, ϕ*_C_(*t*)*, ϕ*_A_(*t*) (third row), their target fractions 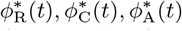 (fourth row), and the instantaneous storing fractions *ϕ*_SC_(*t*) and *ϕ*_SA_(*t*) (solid lines, bottom row)), and their target fractions 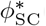 and 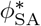 (dashed lines, bottom row). Note that because of the slow proteome relaxation resulting from the slow growth rate λ(*t*) (see Eq. 29), the cell also needs to grow significantly during the night in order to maintain *ϕ*_SA_, necessary to make cyanophycin for growth during the day. This is in marked contrast to the dynamics in the quasi-equilibrium model, in which the proteome relaxes instantly to changing nutrient levels and the cell does not grow at night (see Fig. 1). Please also note that the average growth rate is significantly lower in this slow-proteome model, *〈λ〉*_24_ = 0.039/h, than in the quasi-equilibrium model, *〈λ〉*_24_ = 0.064/h. Other parameter values the same as in Fig. 1: 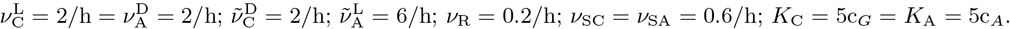

The first point to note is that the average growth rate in the slow-proteome model, *〈λ〉*_24_ = 0.037, is lower than in the quasi-equilibrium model, which is *〈λ〉*_24_ = 0.064. Clearly, the slow relaxation of the proteome drastically lowers the growth rate. The second point is that while the cells predominantly grow during the day (top row), the growth rate during the night is markedly non-zero near the beginning of the night, in marked contrast to the behavior in the quasi-equilibrium model (Fig. 1B),

To characterize the growth dynamics further, we show in the second row of Fig. 3 the concentration of cyanophycin and glycogen, respectively. It is seen that the cyanophycin levels rise during the night, when nitrogen is stored into cyanophycin, yet fall during the day, when the cyanophycin provides the nitrogen source for protein production. Near the end of the day, the cyanophycin levels approach zero, causing the growth rate to drop to zero. The glycogen levels rise during the day, which makes it possible to grow during the night. During the night, however, the glycogen levels rapidly fall, causing the growth at night to come to a halt.

While the behavior of the reservoir dynamics explains the time-dependent growth rate λ(*t*) to a large degree, a few puzzling features remain to be resolved. The first is that the growth rate at the beginning of the day first rises, even though the levels of cyanohycin already fall. The second is that the growth rate drops rather abruptly near the end of the day, even though the concentration of cyanophycin, [N], is well below the enzyme activation threshold *K*_A_. But perhaps the most important question that needs to be addressed is why the cells decide to store glycogen and grow at night, given that the optimal strategy in the quasi-equilibrium model is not to grow at all during the night (see Fig. 1).

To elucidate these questions, we turn to the time traces of the protein fractions, shown in the third to fifth row of Fig. 3. The third row shows *ϕ*_R_*, ϕ*_C_*, ϕ*_A_, while the fourth row shows the target fractions 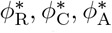 that the cell aims to reach. The fifth row shows the storing fractions *ϕ*_SC_ and *ϕ*_SA_, together with their target fractions, 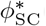 and 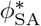, respectively.

To explain the initial rise of the growth rate, we start by noting that at the end of the night, *ϕ*_A_ is large because the cell needs to store cyanophycin during the night, which drains nitrogen flux. The next day, the cell does not need to store nitrogen, while at the beginning of the day the cyanophycin level—the nitrogen source during the day—is still high; taken together this means that the target fraction 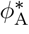 will be relatively low (fourth row). Indeed, at the beginning of the day, the target fraction 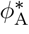 is smaller than the current fraction *ϕ*_A_, causing *ϕ*_A_ to fall initially. This allows *ϕ*_R_ to rise, and since the growth rate is proportional to *ϕ*_R_ (see Eq. 29), this tends to raise the growth rate. The growth thus rises initially, because the proteome slowly adapts to maximize the growth rate.

As time progrresses, the cyanophycin level falls, which causes the target fraction 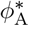 to rise (fourth row). At some point, the current fraction *ϕ*_A_ becomes equal to the target fraction 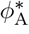. From this moment on, *ϕ*_A_ will rise in order to maintain the flux of nitrogen in the face of the falling cyanophycin levels. This rise in *ϕ*_A_ is accompanied by a drop in *ϕ*_C_ and *ϕ*_R_, causing the growth rate to go down.

Finally, why does the cell grow at night? In this model, the cyanophycin storing proteins are not made during the day, which means that then the storing fraction *ϕ*_SA_ will fall, because of dilution due to growth. Inevitably, at the beginning of the night, the fraction *ϕ*_SA_ will always be smaller than that at the end of the night before. Consequently, *ϕ*_SA_ must rise to move towards the target fraction 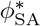, which in this case is close to the maximium at which the growth rate is zero, 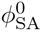 (dashed blue line in last row). However, in the absence of protein degradation, the proteome can only relax because of growth, and, indeed, this is the reason why the cell needs to grow during the night: without growth, *ϕ*_SA_ would eventually become zero, and no cyanophycin could be stored. During the night, new storing proteins have to be made, in order to compensate for the drop in *ϕ*_SA_ resulting from dilution during the day.

Lastly, in order to grow during the night, the cell needs to store glycogen during the day, which explains why *ϕ*_SC_ is non-zero during the day. The cell thus adopts a mixed strategy in which it grows during the day and during the night, because this is the optimal strategy in the presence of slow proteome relaxation. In the next section, we will study whether anticipation makes it possible to counteract the detrimental effects of slow proteome relaxation, by initiating a response ahead of time.

## ANTICIPATION

To study the importance of anticipation, we first consider the scenario where the cell can express the storing fractions *ϕ*_SC_ and *ϕ*_SA_ before the beginning of the day and the night, respectively; here, we thus do not consider the possibility that cells can anticipate the changes in the protein efficiencies 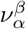 (see Eqs. 41 and 42). More specifically, we consider 4 optimization parameters: the magnitudes of *ϕ*_SC_ and *ϕ*_SA_ and the timings of their expression; to limit the optimisation space, we take the duration of the expression window to be constant, namely 12h. Performing the optimisation, we observed that the growth-rate dependence on the expression timing of *ϕ*_SC_ was rather weak, because, as we will see below, the optimal *ϕ*_SC_ is very small. We therefore considered one anticipation time *T*_a_, which determines the times *k*24 *− T*_a_ and 12 + *k*24 *− T*_a_, with *k* = 0, 1, 2, …, from which *ϕ*_SC_ and *ϕ*_SA_ respectively are expressed for 12 hours at constant values, respectively. This limits the optimisation space to 3 parameters: the magnitudes of *ϕ*_SA_ and *ϕ*_SC_, respectively, and the anticipation time *T*_a_.

To analyze the importance of anticipation, we optimized the growth rate over *ϕ*_SA_ and *ϕ*_SC_ for each value of *T*_a_, for the same set of parameters as in Figs. 1–3. Fig. 4 shows the result. It is seen that expressing the storing enzymes about 4.5 hours before the beginning of the next part of the day can speed up growth by about 20%.

**FIG. 4:**
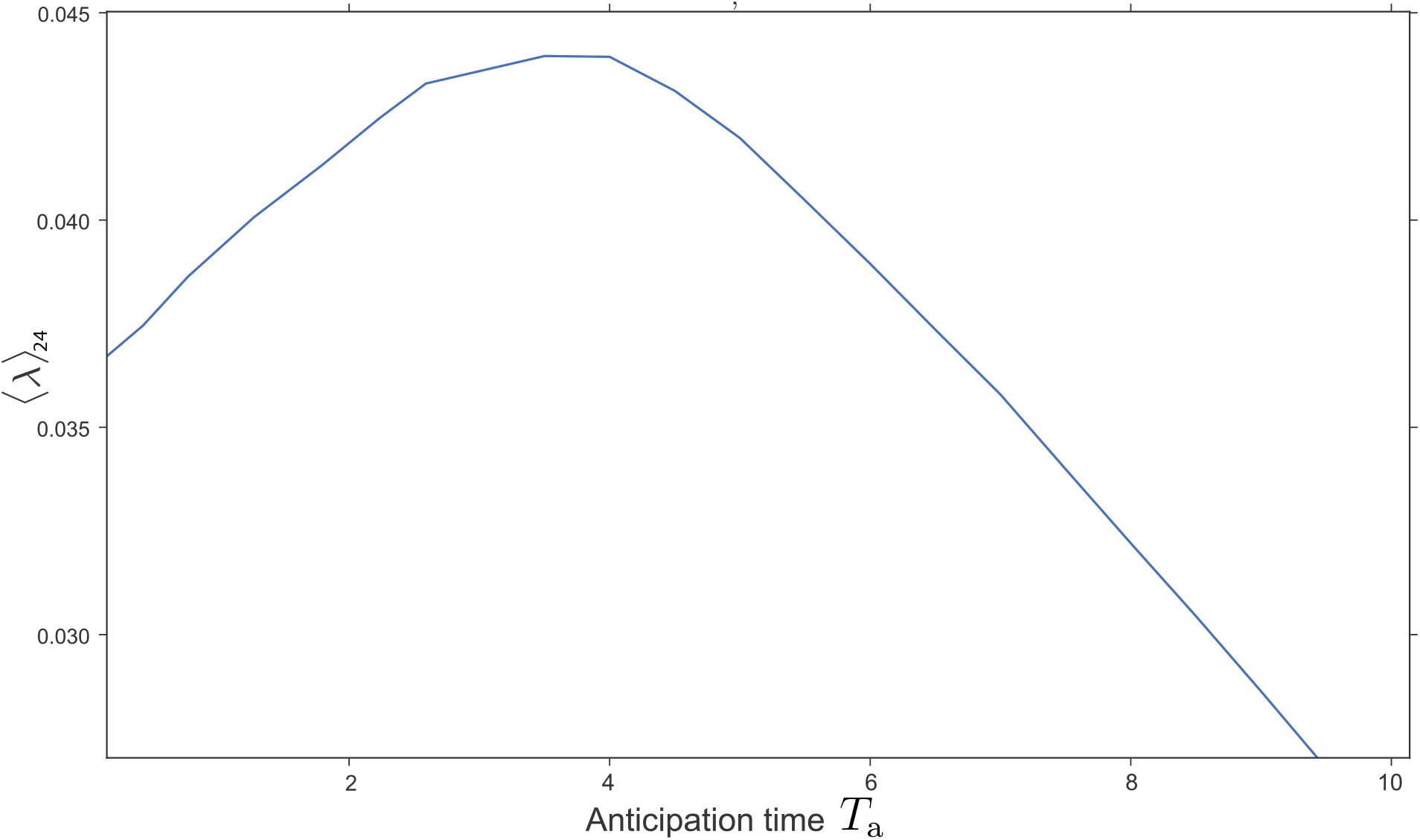
The average growth over 24 hours, *〈λ〉*_24_ (1/h), as a function of the anticipation time *T*_a_ (h) in the full model, including anticipation. The anticipation model is identical to the slow-proteome model in that it is given by Eq. 29 for*λ* (*t*), Eq. 34 for *ϕ_α_* (*t*), and Eqs. 35–40, which are solved to yield 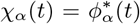 in Eq. 34 and *σ* (*t*) = *σ \** (*t*) in Eq. 29, except that the storing fractions *ϕ*_SC_ and *ϕ*_SA_ start to be expressed an anticipation time *T*_a_ before the beginning of the day and night, respectively; the reservoir dynamics is, as for the other models, given by Eqs. 23 and 24. It is seen that there is an optimal anticipation time 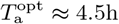 that maximizes the average growth rate over 24 hours. This maximal growth rate is about 20% higher than in the slow-proteome model (see Fig. 3). The principal reason is that the cyanophycins-storing fraction can already be made before the beginning of the night, as elucidated in Fig. 5. Other parameter values the same as in Fig. 1: 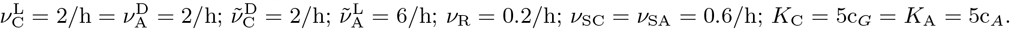

To elucidate this behavior, we show in Fig. 5 the time traces for the optimal anticipation time 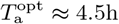 that maximizes the growth rate (Fig. 4). The top row shows that, as in the slow-proteome model, the growth rate first rises at the beginning of the day, because the proteome still adapts to the changing nutrient levels. How ever, at about 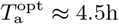 before the end of the day, the growth rate goes down markedly. This is because the cell starts to express the proteins *ϕ*_SA_ that store cyanophycin during the night (bottom row). Clearly, there is a cost to anticipation: it lowers the instantaneous growth rate. The cell should therefore not express the cyanophycin-storing proteins too early in the day. Yet, expressing cyanophycin-storing enzymes already during the day also has a marked benefit: it makes it possible to reach a sufficiently high level of *ϕ*_SA_ before the beginning of the night such that enough cyanophycin can be stored during the night. The cell therefore does not need to grow during the night to raise *ϕ*_SA_, as in the slow-proteome model; indeed, even though *ϕ*_SA_ does not rise during the night, the level is much higher than the average level in the slow-proteome model, so that more cyanophycin is stored during the night, as a result of which the cells grow much faster during the day (compare with Fig. 3). Anticipation thus makes it possible to implement the optimal growth strategy as revealed by the quasi-equilibrium model (see Fig. 1), which is to grow exclusively during the day, as observed experimentally.

**FIG. 5:**
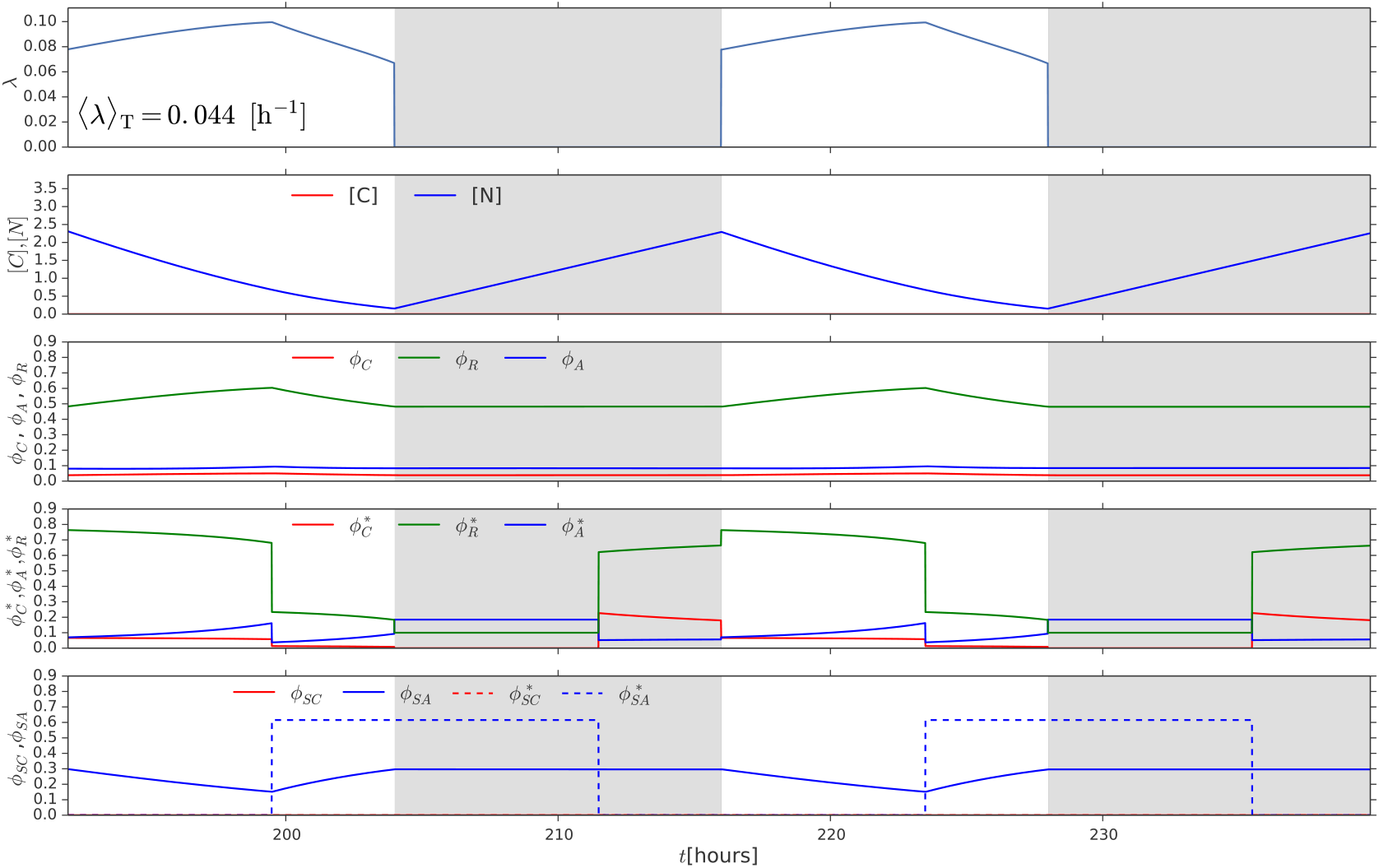
Dynamics of the full model, including anticipation. The model is identical to that of Fig. 4 with the anticipation time *T_a_* equal to its optimal value 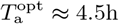. Time traces of the growth rate*λ* (*t*) (1/h, top row), glycogen levels [C](*t*) and cyanophycin levels [N](*t*) (second row), protein fractions *ϕ*_R_(*t*)*, ϕ*_C_(*t*)*, ϕ*_A_(*t*) (third row), their target fractions 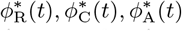 (fourth row), and the instantaneous storing fractions *ϕ*_SC_(*t*) and *ϕ*_SA_(*t*) (solid lines, bottom row)), and their target fractions 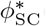(*t*) and 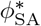(*t*) (dashed lines, bottom row). The average growth rate over 24 hours in this model, *〈λ〉*_24_ = 0.044/h, is about 20% higher than in the slow proteome model. Note also that the growth rate during the day first rises because the proteome is still adapting to the nutrient levels (top row); however, *T*_a_ = 4.5h before the beginning of the night, the growth rate goes down, because the cell prepares for the night by expressing the cyanohycin storing fraction *ϕ*_SA_ (bottom panel). Before the end of the day, *ϕ*_SA_ has reached a level that is sufficient to store enough cyanophycin during the night for fueling growth the next day. Concomitantly, the growth rate is now zero during the night, in contrast to the scenario in the slow-proteome model where *ϕ*_SA_ has to be made during the night, and the cells therefore have to grow during the night Fig. 3. Other parameter values the same as in Fig. 1: 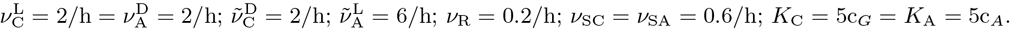

We also considered anticipation of 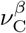 and 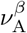, as described around Eqs. 41 and 42. However, because the growth rate is zero at night, the benefit of optimising 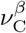, 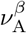 is marginal, for two reasons. Firstly, because the cell cannot grow at night, it cannot adjust the proteome before the beginning of the day. Secondly, adjusting the proteome fractions during the day based on the anticipated efficienies 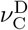 and 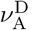 during the night would lower the instantaneous growth rate, because the instantaneous protein fractions *ϕ_α_* would become suboptimal, i.e. not given by the current efficiencies 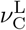 and 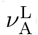.

## DISCUSSION

The power of the framework of Hwa and coworkers is that it provides a coarse-grained description of the proteome with only a limited number of sectors, characterized by enzyme efficiencies that can be measured experimentally [1–5]. We therefore sought to develop a minimal model, consisting of a small number of sectors that can be characterized experimentally, also given the fact that as yet there is no experimental data that warrants a more detailed model. Nonetheless, even though the model consists of only 3 main sectors and 2 storing sectors, the dynamical behavior of our model is already very rich. Specifically, our analysis shows that the requirement to store carbon and nitrogen means that the cells tend to adopt an extreme strategy in which they exclusively grow during the day. The fundamental reason is contained in the growth laws uncovered in refs. [1–5]: storing more glycogen during the day will increase the growth rate during the night, yet this benefit decreases as more cyanophycin-storing enzymes are expressed during the night (and vanishes in fact when this fraction approaches its maximum at which the growth rate becomes zero, see Fig. 2); at the same time, the benefit of storing more cyanophycin during the night—growing faster during the day—increases as less glycogen is stored during the day. The interplay between these two effects creates a positive feedback loop in which the cells store as much cyanophycin as possible during the night and as little glycogen as needed during the day to maximize the growth rate during the day.

Our analysis also reveals that the slow relaxation of the proteome creates a severe challenge in implementing the optimal strategy. In the absence of protein degradation, the cells need to grow in order to adjust their proteome. Yet, cyanobacterial cells grow slowly, which means that the relaxation time will be long compared to the 24h period of the day-night rhythm. In fact, to reach the required cyanophycin-storing fraction, cells would need to grow significantly during the night, in the absence of anticipation. Indeed, the principal benefit of having a circadian clock, according to our model, is that by knowing the time the cyanobacterium can anticipate the shift from day to night and express the storing proteins ahead of time. Interestingly, this prediction appears to be supported by recent mass-spectrometry proteomics data and RNA-sequencing transcriptomics data on *Cyanothece*: the expression of cyanophycin is highest in the late *light* and progressively dimishes during the night into the early light [20].

In a beautiful series of experiments, Johnson and coworkers showed that circadian clocks can provide a fitness benefit to organisms that live in a rhythmic, circadian environment [27, 28]. Mutants of *S. elongatus* with different intrinsic clock periods were competed with wild-type strains and with each other, and the strain whose intrinsic clock period most closely matched that of the light-dark (LD) cycle won the competition [27, 28]. When the intrinsic period of the clock and the period of the LD cycle are altered with respect to one another, then the period of the (driven) clock remains equal to that of the driving signal, as long as the clock remains phase locked to the LD cycle [29]. However, their phase relationship will change [29]. This altered phase relationship is probably the reason why strains with ‘non-resonant’ clock rhythms have a lower growth rate [28]. We investigated whether, according to our model, *Cyanothece* would exhibit similar behavior. To this end, we started from the idea that changing the intrinsic clock period keeping the period of the LD cycle equal to 24h, will alter the phase of the clock. We thus computed the average growth rate *〈λ〉*_24_ as a function of the phase shift *ϕ*. Fig. 6 shows the result. It is seen that changing the phase of the clock from its optimal value can reduce the growth rate by more than 10%. With an incorrectly set clock the cells can no longer accurately anticipate dawn and concomitantly start the production of the cyanophycin storing enzymes either too late or too early.

**FIG. 6:**
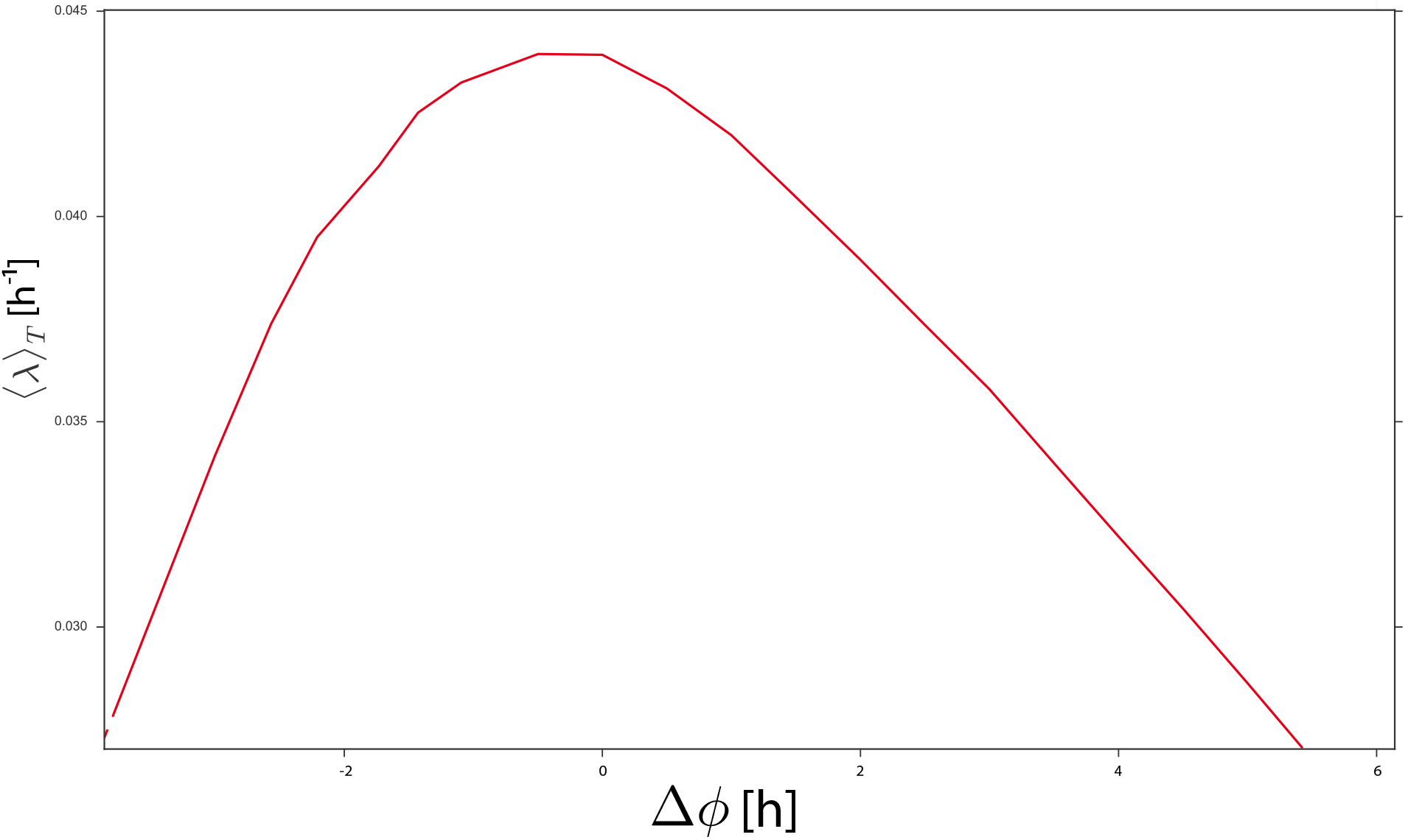
Changing the phase of the clock from its optimal value can significantly reduce the growth rate. The model is identical to that of Fig. 5, meaning that a phase shift Δ*ϕ* = 0 corresponds to the full model with the optimal anticipation time 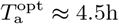. WhenΔ *ϕ* is changed away from 0, all parameters, including the magnitudes of *ϕ*_SC_ and *ϕ*_SA_, are kept constant, except for the time windows in which *ϕ*_SC_ and *ϕ*_SA_ are expressed: these windows are shifted by an amount *ϕ*. Indeed, while in Fig. 4 the magnitudes of *ϕ*_SC_ and *ϕ*_SA_ are optimized for each value of *T_a_*, here the values of *ϕ*_SC_ and *ϕ*_SA_ remain equal to those corresponding to 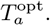 Other parameter values the same as in Fig. 1: 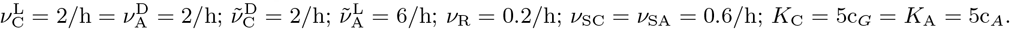

In our model, cyanophycin serves exlucisely as a source of nitrogen, which is a reasonable starting point given that cyanophycin is very rich in nitrogen. However, cyanophycin also contains carbon and it has indeed been speculated that it also provides a carbon store [18]. Our model could be extended to include this. While the benefit of providing a carbon source during the day might be small in the presence of high CO_2_ and light levels, the cost of draining carbon flux at night might be more significant—this effect could be included by adding a term to the equation for the carbon flux (Eq. 3), representing the carbon flux into cyanophycin during the night. Including this effect in the model will also raise the required levels of glycogen.

While the dynamics of our minimal model is already complex, it seems natural to increase the number of sectors as more data becomes available. In particular, it might be of interest to distinguish between proteins of a given sector that are generic, i.e. expressed at signifin-cant levels both during the day and during the night, and proteins that are specific to one part of the day, such as the photosynthesis components. The challenge will be to define major subsectors and devise experiments which make it possible to measure the associated enzyme efficiences.

Another natural extension of our model is to include protein degradation. First of all, active protein degradation makes it possible to increase the proteome relaxation rate. While active protein degradation by itself tends to slow down the growth rate, reaching the optimal proteome partitioning faster might offset this cost. Secondly, some proteins tend to be unstable, meaning that degradation by spontaneous decay is inevitable. In our full model, the amount of glycogen stored is vanishingly small, because our model only considers glycogen as a source of carbon for protein synthesis and the cells do not grow during the night. At the same time, it is well known that cyanobacteria store glycogen. Some of the stored glycogen will be essential for providing the energy to run maintenance processes, such as DNA repair, or to drive the cyanophycin-storing reactions—storing cyanophycin is ATP consuming [31]. However, it is also possible that glycogen is needed to synthesize those proteins that have decayed significantly during the night, such as the components of the protein synthesis machinery. It would then be interesting to see whether including this into the model would yield the prediction that it is beneficial to start expressing these proteins before the end of the night, as observed experimentally [20].

We have focused here on the cyanobacterium *Cyanothece*. However, the application of our framework to cyanobacteria such as *Synechococcus* and *Synechocystis* predicts that also these bacteria tend to grow predominantly during the day (data not shown), as observed experimentally [30]. If the marginal cost of storing glycogen—the reduction in the growth rate during the day—is higher than the marginal benefit—the increase in the growth rate during the night—then the optimal strategy is to not store any glycogen at all for growth during the night, and hence exclusively grow during the day. While this observation may explain why these cyanobacteria predominantly grow during the day, it does not explain why these bacteria have a clock. Indeed, the mechanism by which a clock provides a benefit to *Cyanothece* as predicted by our model—namely that it allows the cell to make storing proteins before it stops growing—does not apply to *Synechococcus* and *Synechocystis*. It is tempting to speculate that the latter cyanobacteria possess a clock because that enables them to replace the photosynthesis proteins which have decayed during the night before the sun rises again.

